# Auxin Regulation and *MdPIN* Expression during Adventitious Root Initiation in Apple Cuttings

**DOI:** 10.1101/2020.03.18.997973

**Authors:** Ling Guan, Yingjun Li, Kaihui Huang, Zong-Ming (Max) Cheng

## Abstract

Adventitious root (AR) formation is critical for the efficient propagation of elite horticultural and forestry crops. Despite decades of research, the cellular processes and molecular mechanisms underlying AR induction in woody plants remains obscure. We examined the details of AR formation in the apple (*Malus domestica*) M.9 rootstock, the most widely used dwarf rootstock for intensive production, and investigated the role of polar auxin transport in post-embryonic organogenesis. AR formation begins with a series of founder cell divisions and elongation of interfascicular cambium adjacent to vascular tissues. This process was associated with a relatively high indole acetic acid (IAA) content and hydrolysis of starch grains. Exogenous auxin treatment promoted cell division, as well as the proliferation and reorganization of the endoplasmic reticulum and Golgi membrane. By contrast, treatment with the auxin transport inhibitor *N*-1-naphthylphthalamic acid (NPA) inhibited cell division in the basal region of the cutting and resulted in abnormal cell divisions during early AR formation. In addition, PIN-FORMED (PIN) transcripts were expressed differentially throughout the whole AR development process, with the up-regulation of *MdPIN8* and *MdPIN10* during induction, an up-regulation of *MdPIN4*, *MdPIN5* and *MdPIN8* during extension, and an up-regulation of all *MdPINs* during AR initiation. This research provides a deeper understanding of the cellular and molecular underpinnings of the AR process in woody plants.

## Introduction

Propagation via cuttings is the most economical method for the mass production of horticultural and forestry plants, while simultaneously maintaining desirable genetic traits (Lei *et al*., 2018a,b). Adventitious root (AR) formation is essential to this process, and its induction has been studied for decades (De Klerk *et al*., 1995; Dubois *et al*., 1988). AR formation is affected by many factors including juvenility, ontology, species/genotypes, and various environmental conditions, such as extreme temperatures, salt stress, and/or the content of H_2_O_2_, NO, and Ca^2+^ (Guan *et al*., 2015), as well as by the endogenous and exogenous application of plant hormones (Uwe *et al*., 2016; Guan *et al*., 2019). Although AR formation has been studied at the anatomical level, its molecular basis, including the mechanism that triggers the process, remains unclear (Bryant *et al*., 2015). AR formation can be divided into three phases: induction, initiation, and extension, which lead to new visible root systems. AR can initiate from internodes, callus formed at the base of cuttings, or the hypocotyl of herbaceous plants (Li *et al*., 2009; Lovell and White, 1986). In apple, ARs emerge from lenticels, in which large intracellular spaces allow for gas exchange in stems.

According to recent anatomical studies in *Arabidopsis thaliana*, founder cell division gives rise to AR primordial (ARP), which then develops into a new root (Della Rovere *et al*., 2013). Similar to that of primary or lateral roots (LRs), the indeterminate growth of AR depends on cell division and elongation, which are affected by various environmental conditions (Nguyen *et al*., 2018). AR formation is regulated by almost all major plant hormones, with auxin strongly promoting the process (Elmongy *et al*., 2018). Exogenous auxin treatment or elevated endogenous auxin levels through genetic engineering increase the rate of AR formation and the number of AR formed, whereas impaired auxin signaling or transport via mutagenesis or auxin transport inhibitors inhibit AR initiation (Dai, *et al*., 2005; Yang *et al*., 2018; Uwe *et al*., 2016).

Auxin gradient acts as a master controller of AR formation, development, and geotropic responses (Park *et al*., 2017; Guan *et al*., 2019). The relationship between the post-embryonic root of AR (and LR) formation and polar auxin transport has been established by using the auxin transport inhibitor 1-*N*-naphthylphthalamic acid (NPA) (Agulló-Antón *et al*., 2014; Liu *et al*., 2009). The promoting effect of polar auxin transport (PAT) on AR development can be counteracted by NPA, which disturbs polar auxin transport and consequently, AR formation (Mignolli *et al*., 2017).

The asymmetric cellular localization of PIN-FORMED (PIN) auxin efflux carriers have been shown to play a rate-limiting role (Weller *et al*., 2017) in directing cell-to-cell auxin flow (Forestan and Varotto, 2012; Zažímalová *et al*., 2007). Despite their importance, the roles of auxin and PINs in regulating this process in woody horticultural plants have rarely been studied. Here, we integrated anatomy and ultrastructural observation, hormone analysis, and the spatiotemporal expression profile of the *PIN* genes to investigate the details of early initiation and organization of AR primordia within the lenticels of (*Malus domestica*) rootstock M.9 cuttings.

## Materials and methods

### Plant material and growth conditions

Six-month-old apple M.9 cuttings were used for all studies. The bases of the cuttings (0-4 cm) were submerged in ventilating hydroponic containers filled with Hoagland’s nutrient solution, pH 5.8 (Eliasson, 1978), and allowed to grow in a growth chamber (~25 °C) under a 16:8 L/D cycle with 300 μmol·m^−2^·s^−1^. All plant samples were collected at 10:00 AM each day and were harvested every 24 hours for 168 hours (7 days). Plant materials used for morphological studies were fixed in 2.5% glutaraldehyde, and those used for phytohormone quantitation and gene expression analysis were frozen in liquid nitrogen and stored at −80 °C until use.

### Treatments and growth analysis

Plants were grown in Hoagland’s nutrient solution (control) or supplemented with 10 μM indole-3-acetic acid (IAA) or N-1-naphthylphthalamic acid (NPA), as positive and negative regulators of polar auxin transport, respectively. IAA and NPA were purchased from Sigma-Aldrich (catalog numbers 12886 and 33371, respectively), and were dissolved in dimethyl sulfoxide.

### Anatomical and ultrastructural observation

Cryosectioning was used to observe anatomical changes during AR initiation. Samples were dehydrated and fixed in 2.5% glutaraldehyde solution according to Chen *et al*. (1985). The division and elongation of AR founder cells were observed with a stereoscopic microscope (Leica MZ 6; Wetzlar, Germany) and ImageJ (v. IJ 1.46r; Schneider et al. 2012) was used for data capture and analysis (Ferreira, 2012). Statistical differences in AR density and percent elongation were determined by ANOVA with Tukey’s post hoc test in SPSS 10 (Meulman, 2000) with *p* <0.05. Cryosectional samples were also used for scanning electron microscopy (SEM) and transmission electron (TEM) microscopy.

### Quantitation of free phytohormones

Submerged apple cuttings with approximately equal weight, length, and diameter for each treatment were collected, and 5-mm segments from the bases of the cuttings were used to determine IAA and zeatin (ZT) contents. The samples were incubated in an ice-cold uptake buffer (1.5% sucrose, 23 mM MES, pH 5.5) for 15 min, followed by two 15-min washes with fresh uptake buffer. The cleared tissue was surface-dried on filter paper and then weighed. IAA and ZT levels were measured using HPLC–ESI–MS/MS; Pan *et al*., 2010). The reverse-phase HPLC gradient parameters and selected reaction monitoring conditions for protonated or deprotonated plant hormones ([M + H] + or [M − H] −) are listed in supplementary Tables S1 and S2. IAA and ZT standards were purchased from Sigma (catalog numbers 12886 and Z0750, respectively).

### RNA extraction and qPCR

RNA was extracted using CTAB, and 1 μL was used as the template for cDNA synthesis using the TaKaRa PrimerScript RT Reagent Kit (RR037B; Liaoning, China). Quantitative PCR (qPCR) was performed with an ABI PRISM 7900HT system (Applied Biosystems; Foster City, CA, USA) using a 10-fold dilution of the cDNA template. The reaction mixture included 5 μL of template, 7.5 μL of SYBR Green PCR master mix (4309155; Applied Biosystems), and l μM of each of the two *MdPIN*-specific primers (Tables S3, S4) in a final volume of 15 μL. Primer efficiency was determined with a standard curve analysis using 5-fold serial dilution of a known amount of template, and amplicon specificity was confirmed by sequencing. The thermal cycle regime consisted of 2 min at 50 °C, 10 min at 95 °C, followed by 40 cycles of 15 sec at 95 °C, 30 sec at 54 °C, and 30 sec at 72 °C. Disassociation curves and gel electrophoresis were used to verify the amplification of a single product. C_T_ values were calculated using the SDS2.1 software (Applied Biosystems) and data were analyzed using the 2^ΔΔCT^ method with *18S rDNA* as a reference gene for normalization (Livak and Schmittgen, 2001).

## Results

### AR initiation in apple cuttings

In order to determine the exact position of AR formation, six-month-old M.9 apple cuttings were cultured in Hoagland’s nutrient solution. The submerged stems were analyzed for AR formation over a seven-day period (168h), during which the lenticels continuously expanded (Fig. 1A, C and F). Noticeable lenticel enlargement was seen at 72h, and new adventitious roots emerging from the lenticels were easily observed as small protrusions at 168h (Fig. 1F). The SEM analysis revealed progressive longitudinal splitting of the submerged lenticels in the controls until a deep fissure was observed at 168h (Fig. 1H, J, M, O, Q and T). IAA treatment accelerated longitudinal splitting in lenticels at each time point. For example, compared with the corresponding control, the lenticel dehiscence became deeper at 72h (Fig. 1K, R) and the lenticel surface was almost completely disrupted at 168h (Fig. 1N, U). By contrast, lenticel longitudinal splitting was very thin and shallow in NPA-treated cuttings at 72h (Fig. 1I, P), and the fissures were still very narrow and shallow at 168h as compared to control cuttings (Fig. 1L, S).

**Fig. 1.**
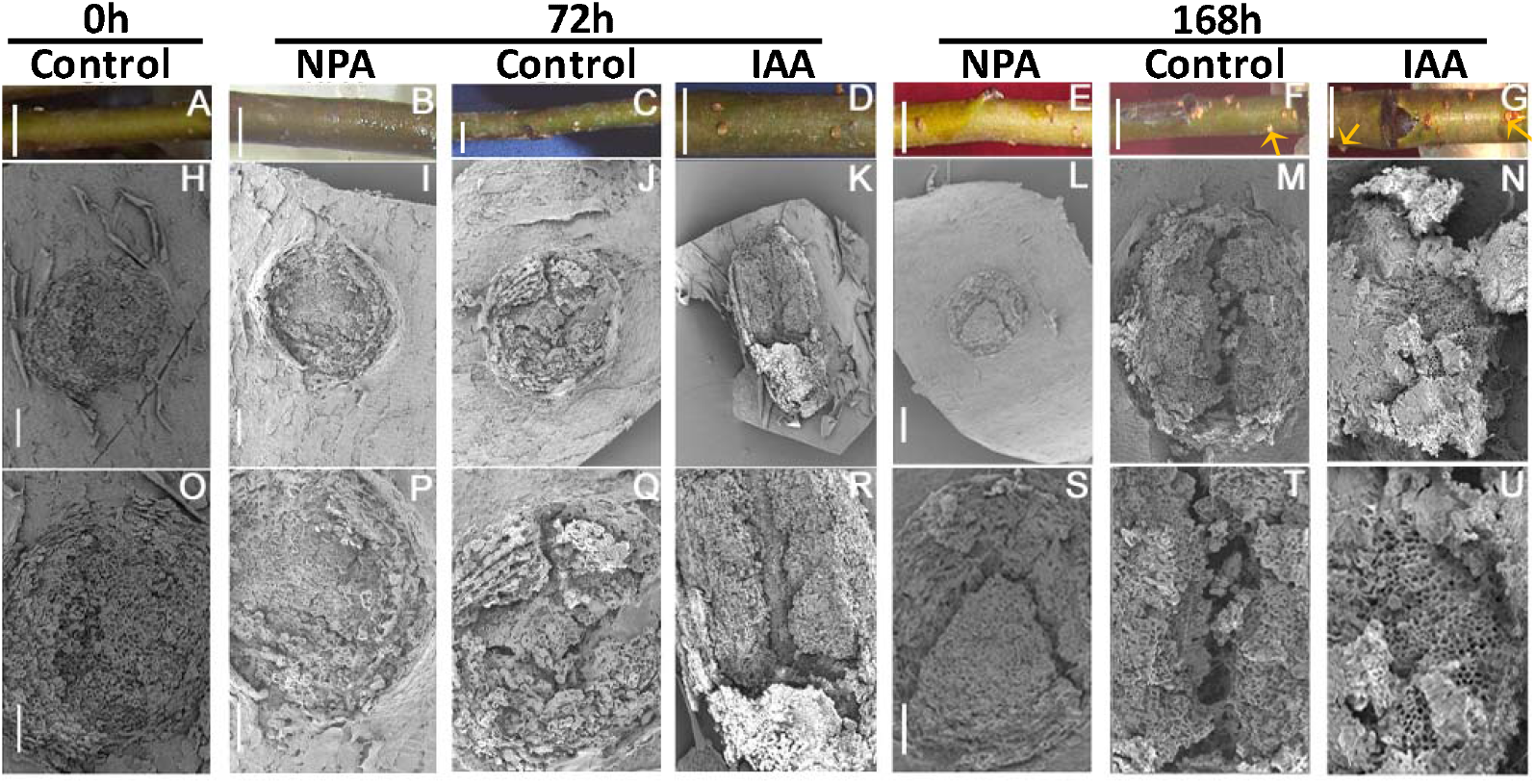
Morphological changes in lenticels upon NPA and IAA treatments. Apple cuttings were cultured in Hoagland’s solution and sampled every 24 hours (h) from 0-168h. Three distinctive development phases were observed at 0, 72, and 168h. A-G: physical appearance of submerged cuttings. Arrows point to the origination of new ARs. H-U, SEM micrographs of lenticels in different AR developmental phases; Bar, A-G=0.4cm, H-U=100 μm, the bar I=J=K; L=M=N; P=Q=R; S=T=U.

The cells underlying the longitudinal splitting in the lenticels were examined to better understand the dehiscence process. The cambial zone in control cuttings had 6–8 layers of cells between the xylem and phloem, and a large number of parenchymatous cells were observed in the interfascicular cambium next to the vascular cylinder (Fig. 2A, E; Fig. S1). During the 72-168h time interval, founder cell divisions produced a large number of cells in the interfascicular cambium adjacent to vascular tissues (Fig. 2A, E, C, G, J, M, P and S). These founder cells exhibited dense cytoplasm and swollen nuclei (Fig. S2), features that are indicative of meristematic activity. Histological observations revealed differences in the founder cells underlying the lenticels in NPA- and IAA-treated cuttings compared with controls. Specifically, NPA-treated lenticels displayed fewer dividing founder cells at 72h and 168h (Fig. 2B, F, I, L, O and R) compared with corresponding controls. By contrast, the lenticels of IAA-treated apple cuttings had more founder and parenchymatous cells at 72 h (Fig. 2B-D, F-H) and more founder cells in the interfascicular cambium at 120h compared with the controls (Fig. 3A), and this process continued through 168h (Fig. 2O-P, R-S). At 168h, the number of elongated founder cells under lenticels continued to increase and started to protrude from the lenticel surface in IAA-treated apple cuttings (Fig. 2Q, T). Together, these results indicate that lenticel dehiscence and AR protrusion begin by 72h and 168h, respectively, and that IAA promotes lenticel longitudinal splitting during early AR emergence.

**Fig. 2.**
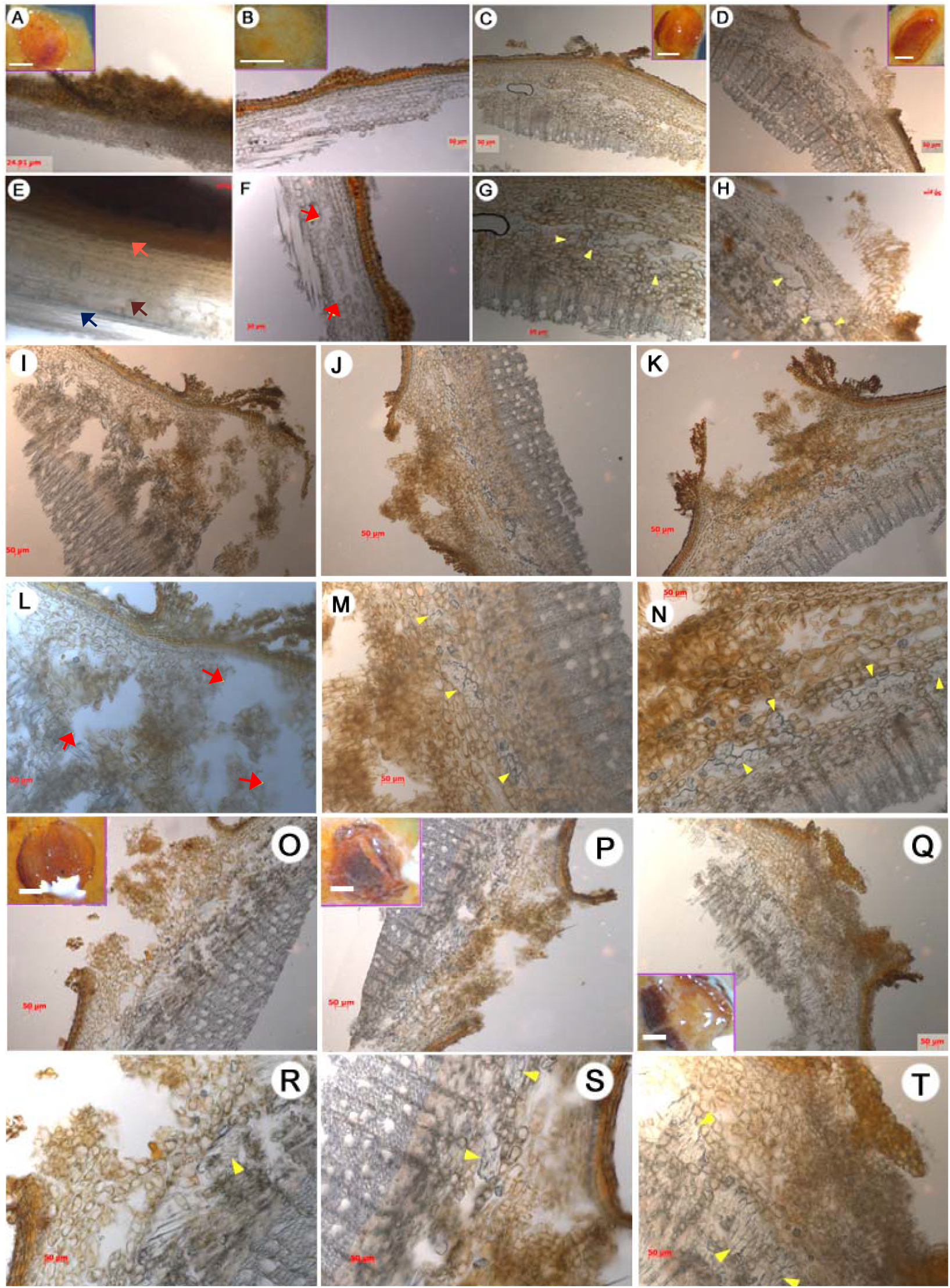
Anatomical observations of lenticels during the rooting of apple cuttings. Each lenticel was observed under the stereomicroscope at 0h (A, E), 72h (B-D, F-H), 120h (I-N), and 168h (O-T). Treatments are: control (A, E, C, G, J, M, P, S), NPA (B, F, I, L, O, R), IAA (D, H, K, N, Q, T). Scale bars, 50 μm. Inset scale bars, A-D=O-Q=0.5 mm. Arrows: yellow, proliferated founder cells; pink, epidermis; brown, parenchyma cells located in interfascicular cambium; dark blue, vascular tissues; red, cavities.

**Fig. 3.**
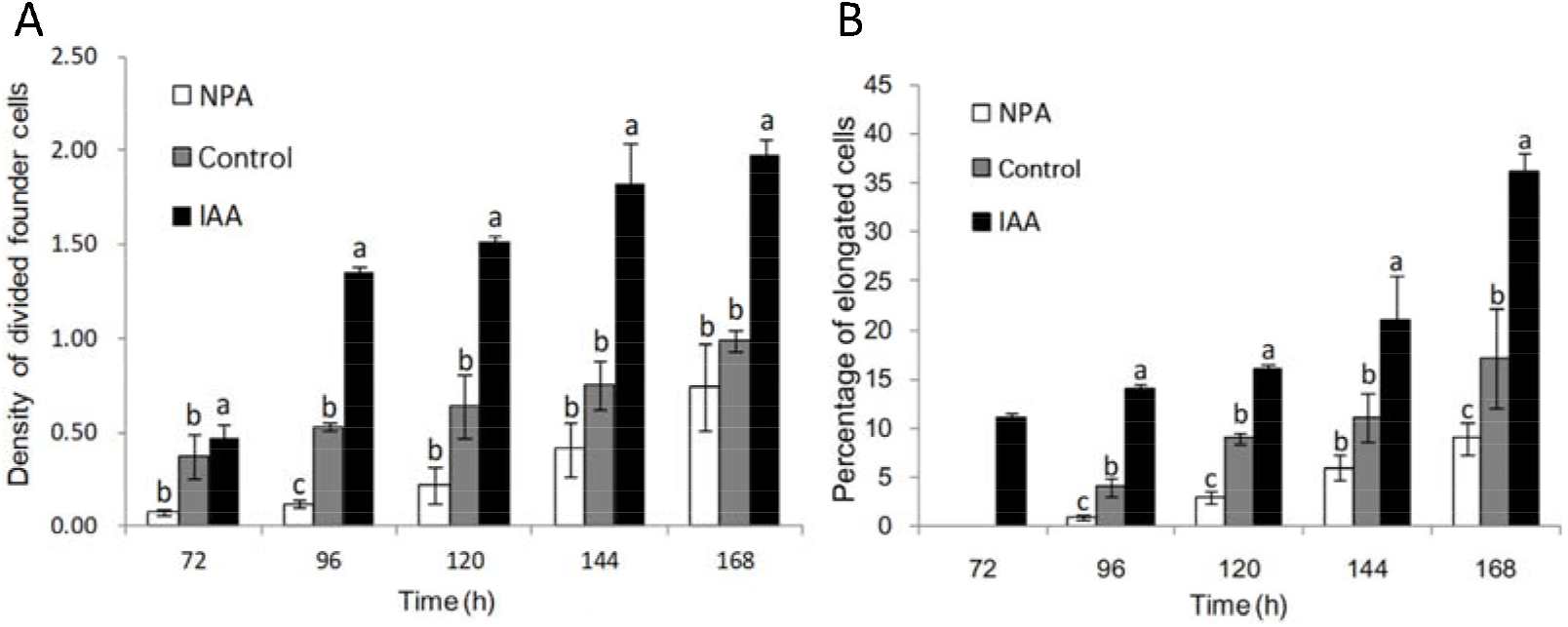
The density and elongation of founder cells during different AR developmental stages. (A). The density of divided founder cells in control, NPA- and IAA-treated apple cuttings. Data are the ratio of divided founder cells to the total number of parenchymal cells. (B). Percentage of elongated cells in control, and NPA- and IAA-treated cuttings. Data are the percentage of the total elongated cells divided by founder cells that have undergone cell division. Elongated cells were at least two times longer than the shorter axis. Error bar represents +/− standard error (SE), n≧15. Bars with different letters indicate statistical significance (*p* < 0.05).

### Auxin controls founder cell division and elongation

By 168h, the density ratio of divided founder cells to total parenchymal cells was two-fold greater in IAA-treated apple cuttings over the control (Fig. 3A). This time point also exhibited the highest number of divided founder cells in the IAA-treated cuttings (Fig. 3A). Taken together, these data suggest that IAA stimulates founder cell division. By contrast, the density of divided founder cells in NPA-treated apple cuttings was approximately half of that of the control cuttings between 72h to 144h (Fig. 3A); further, at 168h, the density of divided founder cells in NPA-treated apple cuttings was 0.75, comparable to that in the control at 144h (Fig. 3A). These observations suggest that NPA delayed founder cell division during late AR initiation. Overall, the density of divided founder cells increased over time in all conditions tested, with IAA and NPA treatments stimulating and inhibiting founder cell division, respectively.

Elongated founder cells were first observed at 72h in IAA-treated cuttings, 24 hours prior to their appearance in the controls (Fig. 3B). Interestingly, elongated founder cells were also observed at 96h in NPA-treated cuttings, but in a lower proportion as compared to the control (Fig. 3B). Like those above, these data support the idea that IAA and NPA stimulate and inhibit founder cell division, respectively. At 168h, elongated founder cells accounted for 16.9%, 35.5%, and 8.8% of total cells in the control, IAA-, and NPA-treated cuttings, respectively. These data are consistent with the changes in divided founder cell density in the respective treatments from 96 to 168h (Fig. 3A), suggesting that AR formation in apple is developmentally programmed.

### Organelle and endomembrane changes during AR initiation

To further investigate the morphological and physiological changes during AR initiation, subcellular structures were examined with transmission electron microscopy (TEM). Numerous starch grains were observed in the plastids at the 0h and 24h time points for all treatments (Fig. 4A-I). However, at 72h, the number and density of starch grains rapidly decreased from the plastids of control and IAA-treated cells, but little change in the density was observed in the plastids of NPA-treated cells (Fig. 4J-L, P-R). At 72h, no changes in starch grain content or organelles were observed in unsubmerged stems (Fig. S3).

**Fig. 4.**
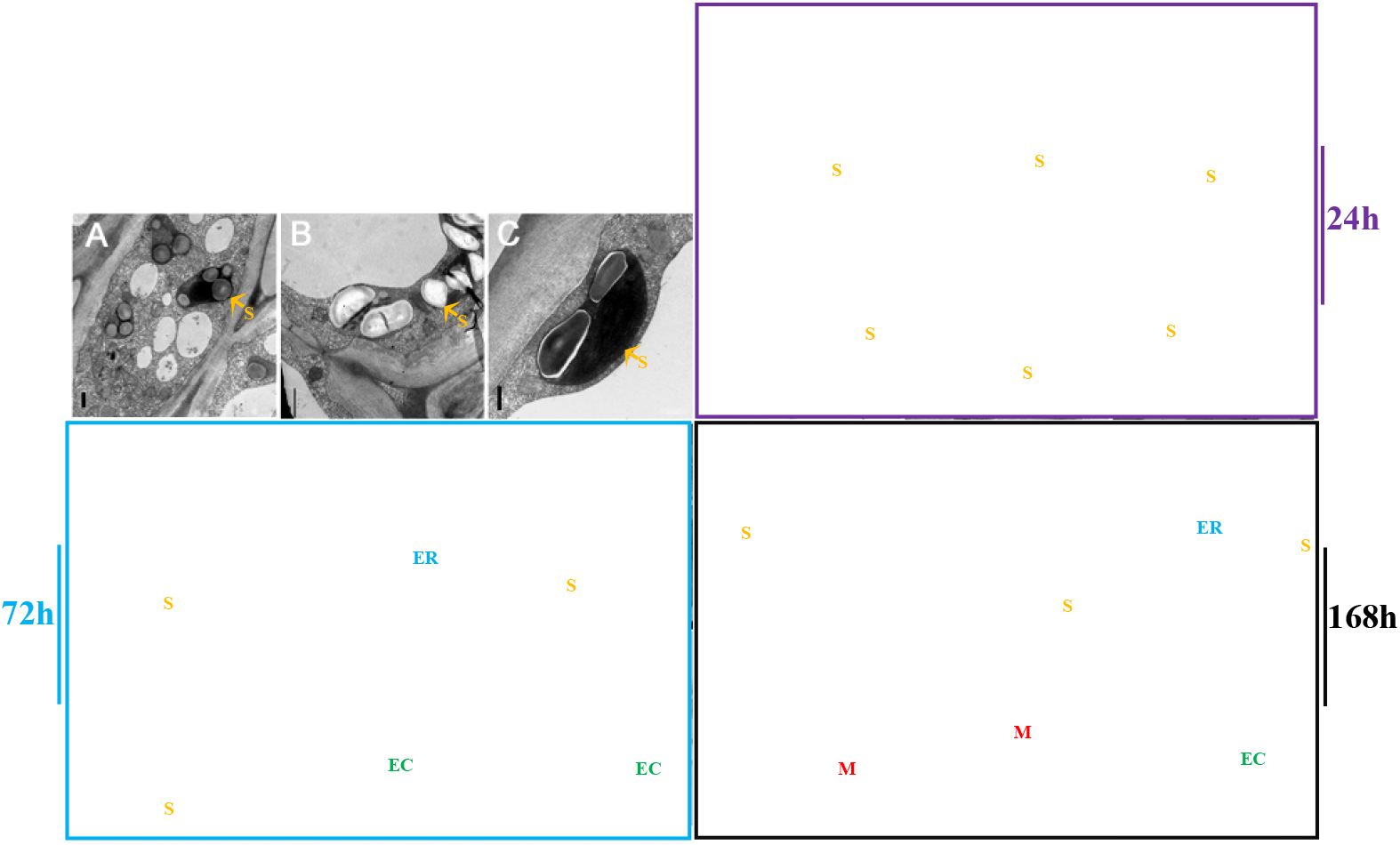
Ultrastructural changes during the rooting of apple cuttings. Transmission electron microscopy of founder cells at 0h (A-C), 24h (D-I), 72h (J-L, P-R), and 168h (M-O, S-U) during AR development. Shown are control (A-C, E, H, K, Q, N, T), NPA treatment (D, G, J, P, M, S), and IAA treatment (F, I, L, R, O, U). Scale bars denote 0.5 μm in A-C, 1.0 μm in N, D, F, G-J, M, P, S and T, 2.5 μm in E and L; 0.25 μm in K and O; 5 μm in Q, R, and U. Orange arrows, starch grains (S); blue arrows, endoplasmic reticulum (ER); green arrows, elongated cells (EC); red arrows, mitochondria (M).

The number and density of the mitochondria, endoplasmic reticulum, and Golgi apparatus in IAA-treated cells were higher than those in control at 72h (Fig. S4). The number of elongated founder cells in control and IAA-treated cuttings continued to increase from 72 (Fig. 4Q-R) to 168h (Fig. 4M-O, S-U). By contrast, many dead cells, and cells with abnormal nuclei, were observed in NPA-treated cuttings (Fig. S4). These observations, taken together with the near constant level of starch grains noted in NPA-treated cuttings, the plausible explanation emerges that NPA inhibits starch grain degradation, thus blocking energy needed for founder cell division and elongation, resulting in abnormal subcellular structures and cell death.

### Auxin and cytokinin contents change in a complementary manner

Auxin and cytokinin are both known to play key roles in AR initiation and elongation (Guan *et al*., 2015). Therefore, we determined the changes in the concentrations of IAA and zeatin (ZT) during AR initiation in control, and IAA- and NPA-treated cuttings (Fig. 5). Although the IAA concentrations differed between treatments, the trends in the changes of IAA levels were consistent between treatments during the test period. For example, IAA concentration increased steadily with time from 48h to120h in all three treatments and peaked at 120h with 1156.4 ng/g FW, 1216.6 ng/g FW, and 572.89 ng/g FW, in control, IAA- and NPA-treatment, respectively (Fig. 5A). At 168h, IAA level in control and NPA-treated cuttings reduced significantly to 707.2 ng/g FW and 338.8 ng/g FW, respectively, but reduced only slightly to 1137.6 ng/g FW in IAA-treated cuttings, compared with those at 120h.

**Fig. 5.**
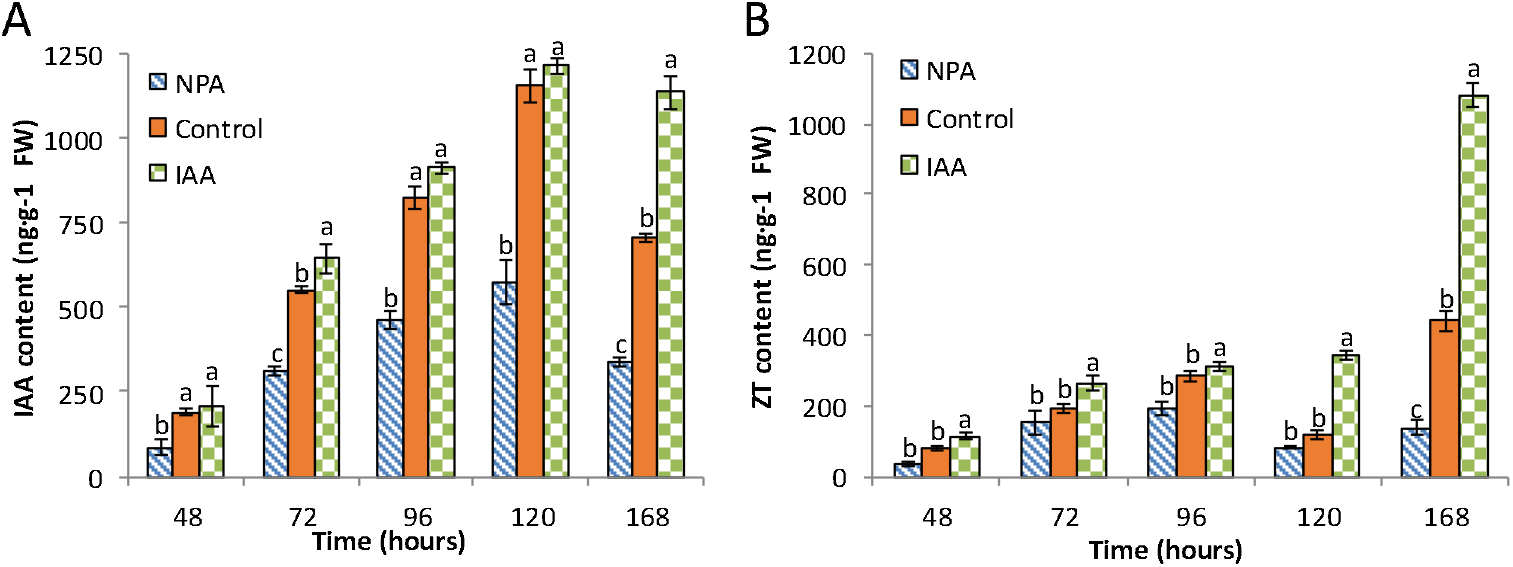
Indole acetic acid (IAA) and zeatin (ZT) levels during different developmental stages of rooting in apple cuttings. Error bar represents +/− SE, n≧3. Different letters indicate significant differences (*p* < 0.05).

In control and NPA-treated cuttings, ZT showed a three-phase accumulation pattern with increasing, then decreasing, and finally increasing again by the end of the experimental period (Fig. 5B). In the control and NPA-treated cuttings, ZT levels increased from 48-96h, decreased at 120h, and increased again at 168h (Fig. 5B). In contrast, this drop was not observed in IAA-treated cuttings, but instead leveled out between time points 96 and 120h, with a significant increase in ZT levels at 120h. Overall, NPA-treated cuttings exhibited lower ZT levels compared to that of the control. For example, ZT levels in NPA-treated cuttings were 36.84% and 72.50% lower than those in control cuttings at 120 and 168h, respectively (Fig. 5B). However, the control and NPA-treated cuttings shared a similar ZT accumulation pattern, which well correlated with that of the change in IAA levels (Fig. 5A). In contrast, ZT levels in IAA-treated cuttings increased with time with no drop in concentration and peaked at 1080.7 ng/g FW at 168h (Fig. 5B), two-fold of that in the control cuttings at 168h.

### MdPIN expression is stage-specific

To correlate *MdPIN* gene expression with the morphological and hormonal changes associated with AR in the apple cuttings, we determined the transcription profiles of the eight *MdPIN* family members at each major stage of AR formation. All eight *MdPIN* gene members were expressed during all three phases of AR formation but exhibited distinct spatiotemporal expression patterns (Fig. 6). For example, *MdPIN1* expression in control cuttings increased from 0h through 96h and peaked at a level 7.2-fold of that at 0h. At 96h, *MdPIN1* expression in control and IAA-treated cuttings was 1.4-fold and 11.2-fold of that at 0h, respectively. In NPA- and IAA-treated cuttings, *MdPIN1* expression showed the same decreasing trend at subsequent time points (Fig. 6A).

**Fig. 6.**
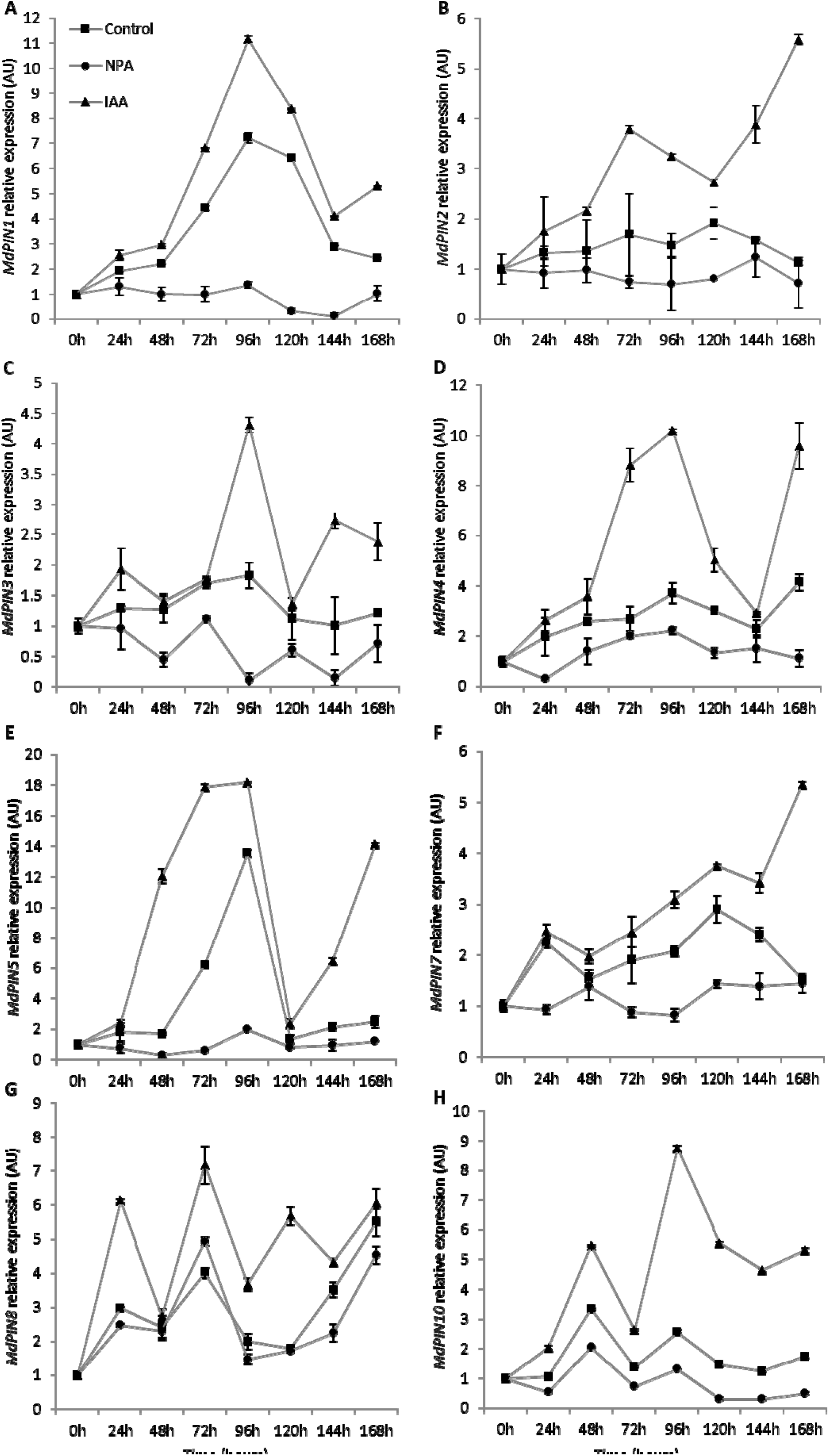
*MdPIN* gene expression. The gene expression levels of *MdPINs* were quantified by qPCR in the bases of apple cuttings during different stages of adventitious root formation. The average relative transcript level of each gene was calculated using the 2^−ΔΔCT^ method; data are shown as mean ± SE (n=3).

*MdPIN2* expression showed relatively uniform expression patterns across the time course, and the fold changes of *MdPIN2* expression between 24h-168h compared with that at 0h ranged from 1.5-2-fold in control and NPA-treated cuttings. *MdPIN2* expression was IAA responsive and increased by up to 5-fold through 24h to 168h (Fig. 6 B). NPA treatment decreased *MdPIN2* expression at all time points.

*MdPIN3* was expressed at relatively low levels during the time course (Fig. 6 C). The expression levels peaked at 96h in control and IAA-treated cuttings and were about 1.8- and 4.3-fold of that at 0h, respectively. In addition, *MdPIN3* expression was significantly inhibited by NPA at all time points, whereas IAA treatment stimulated *MdPIN3* expression, especially from 144h through 168h (Fig. 6 C).

*MdPIN4* expression peaked at 96h and 168h, with an approximately 4-fold increase compared with that at 0h (Fig. 6D). *MdPIN4* expression appeared to be sensitive to IAA and NPA treatments after 48h, with the highest expression levels appearing at 96h 2.2- and 10.2-fold higher than that at 0h in NPA- and IAA-treated cuttings, respectively.

*MdPIN5* had the highest level of expression in the control treatment at 72h and 96h, with a 13.5-fold increase at 96h in comparison of that at 0h (Fig. 6E). *MdPIN5* expression was IAA sensitive and its expression increased in the same pattern observed for controls, with a 18.2-fold increase at 96h. NPA treatment strongly inhibited *MdPIN5* expression in all time points.

*MdPIN7* expression peaked at 24h and 120h, approximately 2.5-3-fold of that at 0h (Fig. 6F). *MdPIN7* expression was sensitive to IAA starting at 72h and expression increased to 5.3-fold at 168h. *MdPIN7* showed little response to NPA treatment.

*MdPIN8* expression increased 3-fold at 24h compared to the baseline at 0h, and IAA treatment induced *MdPIN8* expression during the time course (Fig. 6 G). In contrast, NPA slightly inhibited *MdPIN8* expression. Relatively high expression of *MdPIN8* appeared at 72h and 168h, corresponding to the initiation phase. *MdPIN8* expression peaked at 168h, with 5.5 times of that at 0h. In NPA- and IAA-treated cuttings, *MdPIN8* expression peaked at 72h, with 4.9- and 7.2-fold of that at 0h, respectively. From 96h-168h, NPA treatment increased *MdPIN8* expression.

*MdPIN10* was also expressed during the early stages of AR formation, peaking at 48h, 2-fold above the baseline. At 48h, *MdPIN10* expression in NPA- and IAA-treated cuttings increased by 3.4- and 5.6-fold, respectively, compared with the levels at 0h, (Fig. 6H). In IAA-treated cuttings, *MdPIN10* expression peaked at 8.6-fold at 96h. In contrast to *MdPIN8*, *MdPIN10* expression was inhibited by NPA at all time points.

## Discussion

### AR formation requires the initiation of new founder cells from interfascicular cambium adjacent to vascular tissues

AR formation is crucial for commercial propagation, and depending on the species, there are two mechanisms for AR formation in most woody plants. AR founder cells can either 1) initiate in the stem but remain dormant until the induction of AR formation by environmental conditions (Guan *et al*., 2015), or 2) initiate *de novo* from cells, such as phloem or xylem parenchyma cells, within or adjacent to the vascular tissues, such as cells from interfascicular cambium or the phloem/cambium junction (De Klerk *et al*., 1995; Naija *et al*., 2008; Guan *et al*., 2015). A typical apple stem contains collateral vascular bundles with a ring around the pith (Figs. 1 and 2, Fig. S1). Initially (at 0h), the cells underlying the lenticels in the interfascicular cambium adjacent to vascular tissues had no observable meristematic activity, whereas the founder cells had already undergone numerous cell divisions 72h-168h post-cutting (Fig. 2, Fig. S2). Therefore, AR formation in apple rootstock M.9 cuttings occurs via the second mechanism. This is consistent with that reported recently in forest tree species, especially conifers, that roots are induced from determined or differentiated cells, and further from positions where roots do not normally occur during development (Pizarro *et al*., 2018).

### ARs protrude through lenticels

Lenticel formation culminates with the disintegration of cork cells under the cork cambium that are ultimately replaced by loosely arranged thin-walled cells, also called supplemental cells or filling cells, during early cork layer formation. Later on, epidermis and cork are squeezed until they crack as supplemental cells continue to divide, and eventually split into a series of lip projections, the lenticels (Topa and McLeod, 1986). The tissues inside lenticels provide a natural channel, for not only gas exchange during flooding or other abiotic stress, but also for AR formation and growth. In this study, lenticels of apple cuttings exhibited an increasingly larger crack during early AR formation (Fig.1A-G). Our results showed that at each time point during AR development, the lenticels of control and IAA-treated cuttings were more severely ruptured by founder cell division compared with those of the NPA-treated cuttings (Fig.1H-U). These developmental changes indicate that auxin promotes lenticel dehiscence and AR protrusion from lenticel channels. This process is similar to auxin-mediated channel formation for lateral root (LR) emergence in *Arabidopsis thaliana* (Swarup *et al*., 2008; Stoeckle *et al*., 2018). In both cases, auxin uptake into cortical and epidermal cells overlying the LR primordium during emergence results in increased expression of genes encoding cell wall-remodeling enzymes, such as pectate lyase, to promote cell separation for LR emergence (Swarup *et al*., 2008; Stoeckle *et al*., 2018). In addition, histological observations detected intra-stem channels formed by enlarged cavities in the interfascicular cambium, and the number and dimensions of these cavities increased over time (Fig. 2). These results further suggest that the cells located in front of newly formed founder cells might have undergone programmed cell death (PCD) to allow AR primordium development, supporting the conclusion of a previous study on AR formation in tomato (Steffens and Sauter, 2005).

### The induction phase of AR formation in apple cuttings

AR formation in apple cuttings can be divided into three phases based on characteristic cellular changes as we described in the anatomical observations: induction (0-72h), initiation (72-120h), and extension (120-168h). In control and IAA-treated cuttings, the induction phase at 72h was characterized by an increased number of founder cells with dense cytoplasm and swallow nuclei (Fig.2 G-H), as well as the hydrolysis of starch grains (Fig. 3 P-Q), which presumably provided energy for the further division and elongation of AR founder cells. By contrast, a large number of starch grains remained in the chloroplasts at 72h in NPA-treated apple cuttings (Fig.4 J), suggesting that NPA had inhibited starch hydrolysis during AR formation whereas IAA enhanced the conversion of starch into “cash energy” for AR induction. These findings suggest that IAA modulates AR induction by inducing the dedifferentiation of interfascicular cambium cells into AR founder cells.

Therefore, AR initiation begins with the division and elongation of a large number of clustered founder cells that fill the cavity opened by lenticel splitting in control and IAA-treated cuttings (Fig.2M-N; Fig. 3F-N), whereas founder cell division was severely impaired in NPA-treated apple cuttings (Fig.2 L). In addition, IAA treatment significantly increased the ratio of divided founder cells to total parenchymal cells, and founder cell density increased more quickly than in controls (Fig. 3A). These data demonstrate the promoting effect of IAA on primordial founder cell division upon the induction of AR formation (Fig. 3A).

With elongation of the founder cells, AR formation advances into the extension phase when the percentage of elongated cells among divided founder cells increased (Fig. 4P-Q, S-T). Elongated AR founder cells appeared earlier in IAA-treated cuttings and were almost twice as abundant as those in control cuttings (Fig. 3B). Thus, we speculate that both lenticel dehiscence and intra-stem PCD (Steffens and Sauter, 2005) might regulate AR formation by modulating polar auxin transport, which regulates auxin gradient.

### Starch grain depletion is associated with AR initiation

Previous studies have found that maintaining an appropriate auxin level and gradient in the basal portion of shoots is essential for AR formation (Kitomi *et al*., 2008, Agulló-Antón *et al*., 2014; Xu *et al*., 2017) but the exact mechanism underlying this is still unclear. Here, we observed rapid starch grain depletion upon the initiation of AR formation in apple cuttings. It has been proposed that starch accumulation and depletion may be a biochemical indicator for early root formation in hypocotyl cuttings, which provides energy for AR formation (references). This energy supplementation process is positively regulated by auxin (Jásik *et al*., 1997; Li *et al*., 2000; Husen *et al*., 2007; Husen *et al*., 2017). In agreement with the published data, our temporal and spatial analysis study also supports a key role of starch grain accumulation and degradation in AR of woody plants, with IAA exerting a stimulatory effect on AR formation and NPA delaying the process (Fig. 4). In addition, starch grain depletion was also associated with endomembrane proliferation and reorganization (Fig. 4 and Fig. S4). Although the physiological basis of how the two processes relate to AR formation, the fact that grain depletion and new membrane system formation were promoted by IAA and inhibited by NPA suggests that polar auxin transport is the initial driving force for these processes (Yu *et al*., 2016).

### Roles of auxin and cytokinin during AR formation

Recent studies have shown that ZT can suppress adventitious root primordium formation whereas auxin signaling positively regulates this complex biological process, suggesting that relatively high IAA and low ZT levels are prerequisites for AR induction and initiation (Li *et al*., 2018; Mao *et al*., 2018). Cytokinins and auxin seem to exert opposite effects on AR formation (Dubois *et al*., 1988; Yang *et al*., 2017). In addition, auxin can directly down-regulate cytokinin biosynthesis while cytokinins have little effect on auxin biosynthesis (Nordström *et al*., 2004). The initial high levels of auxin in cuttings at the induction and initiation phases of AR formation were replaced by high levels of cytokinin in the extension phase, suggesting a crosstalk between auxin and cytokinin metabolism, although the molecular basis remains to be characterized.

Auxin appears to regulate AR formation mainly by modulating AR founder cell division and elongation during the induction and initiation stages (Fig. 2, 3). Similar to that had been observed for LR formation in *Arabidopsis thaliana* (Petersson *et al*., 2009), IAA levels in apple cuttings continued to increase during the induction (0-72h) and initiation (72-120h) phases, maximizing at 120h, followed by a steady reduction after entering the elongation phase (120-168h). ZT levels also increased during the induction and initiation phases and peaked at 96h before they started to decrease. By contrast, NPA hampered IAA accumulation but had little effect on ZT levels. These results point to the maintenance of auxin and cytokinin homeostasis via feedback regulation.

### Correlations between MdPIN expression and AR formation

Most *PINs* function in directional auxin transport while displaying differential expression patterns, reflecting their potential multifaceted roles in plant development (He *et al*., 2017; Short *et al*.; 2018; Wang *et al*., 2017). In Arabidopsis, members of the *PIN* gene family are shown to participate in vascular polar auxin transport, root patterning, root gravitropism, sink-driven auxin gradient establishment, and apical-basal polarity (Friml *et al*., 2002a; Friml *et al*., 2002b; Friml *et al*., 2003; Gälweiler *et al*., 1998; Mravec *et al*., 2009; Muller *et al*., 1998). The *PIN* family in apple has eight members, all of which are expressed differntially during AR formation (Fig. 6). Therefore, we proposed a possible model of the regulatory mechanism to map the main function of each *MdPIN* during the rooting of apple cuttings (Fig. 7A).

**Fig. 7.**
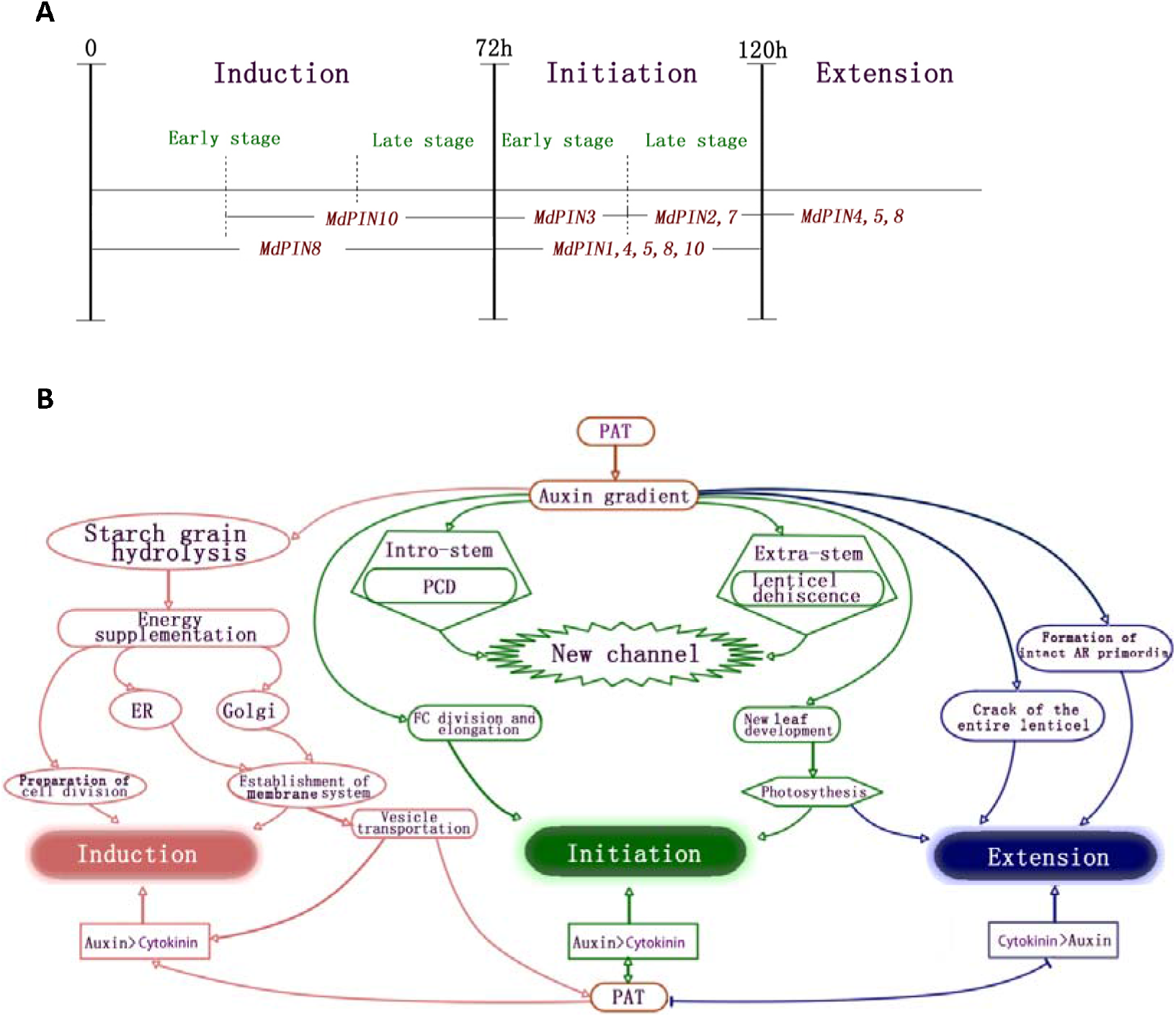
Stage-specific *MdPIN* gene expression patterns and a model of cellular regulatory mechanism during AR formation. (A) Differential expression of *MdPIN*s during AR induction, initiation and extension. Dashed lines represent 24h time points. *MdPIN8* expression peaked during the induction phase (24h), indicating that *MdPIN8* may play a major role in the early stage of AR induction. *MdPIN10* expression peaked at 48h, and was mostly low at other time points, suggesting that *MdPIN10* also contributes to the induction and initiation of AR. The maximum expression of *MdPIN1*, *MdPIN3*, *MdPIN4*, *MdPIN5* occurred at 96h, just 24h prior to the morphological changes in the initiation phase, which ends at 120 h. Further, *MdPIN1*, *MdPIN2*, *MdPIN4*, and *MdPIN5* expression remained high at 120h, indicating that all of these genes also function during the late stage of initiation. *MdPIN7* expression peaked in 120h, suggesting that *MdPIN7* may mainly function in the late stage of initiation. At 168h, differential increases in *MdPIN4*, *MdPIN5*, and *MdPIN8* expression were observed, suggesting that these members promote lenticel dehiscence and AR protrusion from the epidermis in the extension phase.

Up-regulation of *MdPIN8* and *MdPIN10* was mainly associated in AR induction. The initiation phase was associated with an up-regulation of *MdPIN1*, *MdPIN4*, *MdPIN5*, *MdPIN8*, and *MdPIN10*; *MdPIN3* is up-regulated at the early stage of initiation, and *MdPIN2* and *MdPIN7* were up-regulated toward the very end of the initiation stage (Fig. 7A). The extension phase is mainly associated with an up-regulation of *MdPIN4*, *MdPIN5*, and *MdPIN8* (Fig. 7A). Our data suggest that the eight *MdPINs* likely play different roles throughout the AR process, where their gene products may directly or indirectly regulate AR formation in a cooperative manner by mediating anatomical and physiological changes in the cuttings (Fig. 7B). Future biochemical studies should focus on the diverse expression patterns of *MdPIN* proteins to further investigate the function of each in regulating this post-embryonic rooting process.

## Conclusions

AR formation in apple is a coordinated developmental process. IAA stimulates the process likely via the up-regulation of *PIN* gene expression. Auxin stimulates AR founder cells division and elongation probably by promoting energy supplementation via starch grain hydrolysis, which leads to endomembrane system proliferation, lenticel dehiscence, and AR emergence in the apple M.9 rootstock.

## Acknowledgements

We sincerely thank Yi Li from University of Connecticut and Dr. Angus Murphy, Wendy Peer and Jun Zhang from University of Maryland for their valuable suggestions. We thank Jun Hu, Jin Wang, and Zuli Gu for helping with the experimental material and data collection in this study. This work was partially supported by the National Natural Science Foundation of China (Grant no. 31601738).

## References

Agulló-Antón MÁ, Ferrández-Ayela A, Fernández-García N, Nicolás C, Albacete A, Pérez-Alfocea F, Sánchez-Bravo J, Pérez-Pérez JM, Acosta M. 2014. Early steps of adventitious rooting: morphology, hormonal profiling and carbohydrate turnover in carnation stem cuttings. Physiologia Plantarum 150, 446–462.

Brender E, Burke A, Glass RM. 2005. Frozen section biopsy. JAMA 294, 3200–3200.

Bryant PH, Trueman SJ. 2015. Stem anatomy and adventitious root formation in cuttings of Angophora, Corymbia and Eucalyptus. Forests 6, 1227–1238.

Chen T-H, Lam L, Chen S-C. 1985. Somatic embryogenesis and plant regeneration from cultured young inflorescences of *Oryza sativa* L.(rice). Plant cell, Tissue and Organ culture 4, 51–54.

Dawood T, Yang X, Visser EJW, Te Beek TAH, Kensche PR, Cristescu SM, Lee S, Floková K, Nguyen D, Mariani C. 2016. A co-opted hormonal cascade activates dormant adventitious root primordia upon flooding in *Solanum dulcamara*. Plant Physiology 170, 2351–2364.

Della Rovere F, Fattorini L, D’Angeli S, Veloccia A, Falasca G, Altamura MM. 2013. Auxin and cytokinin control formation of the quiescent centre in the adventitious root apex of *Arabidopsis*. Annals of Botany 112, 1395–1407.

Dubois LAM, De Vries DP. The effect of cytokinin and auxin on the sprouting and rooting and rooting of ‘amada’ rose softwood cuttings. Acta Horticulturae 226, 455–464.

Eliasson L. 1978. Effects of nutrients and light on growth and root formation in *Pisum sativum* cuttings. Physiologia Plantarum 43, 13–18.

Elmongy MS, Cao Y, Zhou H, Xia Y. 2018. Root development enhanced by using indole-3-butyric acid and naphthalene acetic acid and associated biochemical changes of in vitro Azalea microshoots. Journal of Plant Growth Regulation 37, 813–825.

Fisher J, Gaillard P, Fellbaum CR, Subramanian S, Smith S. 2018. Quantitative 3D imaging of cell level auxin and cytokinin response ratios in soybean roots and nodules. Plant, Cell & Environment 41, 2080–2092.

Forestan C, Varotto S. 2012. The role of PIN auxin efflux carriers in polar auxin transport and accumulation and their effect on shaping maize development. Molecular Plant 5, 787–798.

Friml J, Benková E, Blilou I, Wisniewska J, Hamann T, Ljung K, Woody S, Sandberg G, Scheres B, Jürgens G. 2002a. *AtPIN4* Mediates Sink-Driven Auxin Gradients and Root Patterning in *Arabidopsi*. Cell 108, 661–673.

Friml J, Vieten A, Sauer M, Weijers D, Schwarz H, Hamann T, Offringa R, Jürgens G. 2003. Efflux-dependent auxin gradients establish the apical–basal axis of Arabidopsis. Nature 426, 147–153.

Friml J, Wiśniewska J, Benková E, Mendgen K, Palme K. 2002b. Lateral relocation of auxin efflux regulator PIN3 mediates tropism in *Arabidopsis*. Nature 415, 806–809.

Klerk G-JD, Keppel M, Brugge JT, Meekes H. 1995. Timing of the phases in adventitous root formation in apple microcuttings. Journal of Experimental Botony 46, 965–972.

Gälweiler L, Guan C, Müller A, Wisman E, Mendgen K, Yephremov A, Palme K. 1998. Regulation of polar auxin transport by *AtPIN1* in *Arabidopsis* vascular tissue. Science 282, 2226–2230.

Guan L, Murphy AS, Peer WA, Gan L, Li Y, Cheng Z-M. 2015. Physiological and Molecular Regulation of Adventitious Root Formation. Critical Reviews in Plant Sciences 34, 506–521.

Guan L, Tayengwa R, Cheng ZM, Peer WA, Murphy AS, Zhao M. 2019. Auxin regulates adventitious root formation in tomato cuttings. BMC Plant Biology 19, 435.

He P, Zhao P, Wang L, Zhang Y, Wang X, Xiao H, Yu J, Xiao G. 2017. The PIN gene family in cotton (*Gossypium hirsutum*): genome-wide identification and gene expression analyses during root development and abiotic stress responses. BMC Genomics 18, 507.

Husen A, Iqbal M, Siddiqui SN, Sohrab SS, Masresha G. 2017. Effect of indole-3-butyric acid on clonal propagation of mulberry (*Morus alba* L.) stem cuttings: rooting and associated biochemical changes. Proceedings of the National Academy of Sciences, India Section B: Biological Sciences 87, 161–166.

Husen A, Pal M. 2007. Metabolic changes during adventitious root primordium development in *Tectona grandis* Linn. f.(teak) cuttings as affected by age of donor plants and auxin (IBA and NAA) treatment. New Forests 33, 309–323.

Jasik J, De Klerk GJ. 1997. Anatomical and ultrastructural examination of adventitious root formation in stem slices of apple. Biologia Plantarum 39, 79–90.

Kitomi Y, Ogawa A, Kitano H, Inukai Y. 2008. *CRL4* regulates crown root formation through auxin transport in rice. Plant Root 2, 19–28.

Klerk G-JD, Keppel M, Brugge JT, Meekes H. 1995. Timing of the phases in adventitous root formation in apple microcuttings. J Exp Bot 46, 965–972.

Lei C, Fan S, Li K, Meng Y, Mao J, Han M, Zhao C, Bao L, Zhang D. 2018. iTRAQ-based proteomic analysis reveals potential regulation networks of IBA-induced adventitious root formation in apple. International journal of molecular sciences 19, 667.

Li F, Sun C, Li X, Yu X, Luo C, Shen Y, Qu S. 2018a. The effect of graphene oxide on adventitious root formation and growth in apple. Plant Physiology and Biochemistry 129, 122–129.

Li K, Liang Y, Xing L, Mao J, Liu Z, Dong F, Meng Y, Han M, Zhao C, Bao L. 2018b. Transcriptome analysis reveals multiple hormones, wounding and sugar signaling pathways mediate adventitious root formation in apple rootstock. International journal of molecular sciences 19, 2201.

Li M, Leung DWM. 2000. Starch accumulation is associated with adventitious root formation in hypocotyl cuttings of *Pinus radiata*. Journal of Plant Growth Regulation 19.

Li S-W, Xue L, Xu S, Feng H, An L. 2009. Mediators, genes and signaling in adventitious rooting. The Botanical Review 75, 230–247.

Liu S, Wang J, Wang L, Wang X, Xue Y, Wu P, Shou H. 2009. Adventitious root formation in rice requires *OsGNOM1* and is mediated by the *OsPINs* family. Cell research 19, 1110–1119.

Livak KJ, Schmittgen TD. 2001. Analysis of relative gene expression data using real-time quantitative PCR and the 2^−ΔΔCT^ method. Methods 25, 402–408.

Márquez G, Alarcón MV, Salguero J. 2019. Cytokinin inhibits lateral root development at the earliest stages of lateral root primordium initiation in maize primary root. Journal of Plant Growth Regulation 38, 83–92.

Mao J, Zhang D, Meng Y, Li K, Wang H, Han M. 2019. Inhibition of adventitious root development in apple rootstocks by cytokinin is based on its suppression of adventitious root primordia formation. Physiologia Plantarum 166, 663–676.

Meulman JJ, Heiser WJ. 1999. SPSS Categories 10.0: SPSS Incorporated.

Mignolli F, Mariotti L, Picciarelli P, Vidoz ML. 2017. Differential auxin transport and accumulation in the stem base lead to profuse adventitious root primordia formation in the aerial roots (*aer*) mutant of tomato (*Solanum lycopersicum* L.). Journal of Plant Physiology 213, 55–65.

Mravec J, Skupa P, Bailly A, Hoyerova K, Krecek P, Bielach A. 2009. Subcellular homeostasis of phytohormone auxin is mediated by the ER-localized PIN5 transporter. Nature 459.

Muller A, Guan C, Galweiler L, Tanzler P, Huijser P, Marchant A, Parry G, Bennett M, Wisman E, Palme K. 1998. *AtPIN2* defines a locus of Arabidopsis for root gravitropism control. EMBO J 17, 6903–6911.

Naija S, Elloumi N, Jbir N, Ammar S, Kevers C. 2008. Anatomical and biochemical changes during adventitious rooting of apple rootstocks MM 106 cultured in vitro. Comptes rendus biologies 331, 518–525.

Nguyen T-N, Tuan PA, Mukherjee S, Son S, Ayele BT. 2018. Hormonal regulation in adventitious roots and during their emergence under waterlogged conditions in wheat. Journal of Experimental Botany 69, 4065–4082.

Nordström A, Tarkowski P, Tarkowska D, Norbaek R, Åstot C, Dolezal K, Sandberg G. 2004. Auxin regulation of cytokinin biosynthesis in *Arabidopsis thaliana*: a factor of potential importance for auxin–cytokinin-regulated development. Proceedings of the National Academy of Sciences 101, 8039–8044.

Pan X, Welti R, Wang X. 2010. Quantitative analysis of major plant hormones in crude plant extracts by high-performance liquid chromatography–mass spectrometry. Nature protocols 5, 986.

Park S-H, Elhiti M, Wang H, Xu A, Brown D, Wang A. 2017. Adventitious root formation of in vitro peach shoots is regulated by auxin and ethylene. Scientia Horticulturae 226, 250–260.

Petersson SV, Johansson AI, Kowalczyk M, Makoveychuk A, Wang JY, Moritz T, Grebe M, Benfey PN, Sandberg G, Ljung K. 2009. An auxin gradient and maximum in the Arabidopsis root apex shown by high-resolution cell-specific analysis of IAA distribution and synthesis. Plant Cell 21, 1659–1668.

Druege U, Franken P, Hajirezaei MR. 2016. Plant hormone homeostasis, signaling, and function during adventitious root formation in cuttings. Front Plant Sci 7:381.

Sánchez-García AB, Ibáñez S, Cano A, Acosta M, Pérez-Pérez JM. 2018. A comprehensive phylogeny of auxin homeostasis genes involved in adventitious root formation in carnation stem cuttings. PloS One 13.

Sauter M. 2005. Epidermal cell death in rice is regulated by ethylene, gibberellin, and abscisic acid. Plant Physiology 139, 713–721.

Schneider CA, Rasband WS, Eliceiri KW. 2012. NIH image to ImageJ: 25 years of image analysis. Nature Methods 9, 671–675.

Short E, Leighton M, Imriz G, Liu D, Cope-Selby N, Hetherington F, Smertenko A, Hussey PJ, Topping JF, Lindsey K. 2018. Epidermal expression of a sterol biosynthesis gene regulates root growth by a non-cell-autonomous mechanism in *Arabidopsis*. Development 145, dev160572.

Stoeckle D, Thellmann M, Vermeer JEM. 2018. Breakout—lateral root emergence in *Arabidopsis thaliana*. Curr Opin Plant Biol 41, 67–72.

Swarup K, Benková E, Swarup R, et al. 2008. The auxin influx carrier LAX3 promotes lateral root emergence. Nature cell biology 10, 946–954.

Topa MA, McLeod KW. 1986. Aerenchyma and lenticel formation in pine seedlings: a possible avoidance mechanism to anaerobic growth conditions. Physiologia Plantarum 68, 540–550.

van Berkel K, de Boer RJ, Scheres B, ten Tusscher K. 2013. Polar auxin transport: models and mechanisms. Development 140, 2253–2268.

Wang L, Guo M, Li Y, Ruan W, Mo X, Wu Z, Sturrock CJ, Yu H, Lu C, Peng J. 2018. LARGE ROOT ANGLE1, encoding OsPIN2, is involved in root system architecture in rice. Journal of Experimental Botany 69, 385–397.

Weller B, Zourelidou M, Frank L, Barbosa ICR, Fastner A, Richter S, Jürgens G, Hammes UZ, Schwechheimer C. 2017. Dynamic PIN-FORMED auxin efflux carrier phosphorylation at the plasma membrane controls auxin efflux-dependent growth. Proceedings of the National Academy of Sciences 114, E887–E896.

Xu D, Miao J, Yumoto E, Yokota T, Asahina M, Watahiki M. 2017a. *YUCCA9*-mediated auxin biosynthesis and polar auxin transport synergistically regulate regeneration of root systems following root cutting. Plant and Cell Physiology 58, 1710–1723.

Xu X, Li X, Hu X, Wu T, Wang Y, Xu X, Zhang X, Han Z. 2017b. High miR156 expression is required for auxin-induced adventitious root formation via *MxSPL26* independent of *PIN*s and *ARF*s in *Malus* xiaojinensis. Front Plant Sci 8, 1059.

Yang X, Jansen MJ, Zhang Q, Sergeeva L, Ligterink W, Mariani C, Rieu I, Visser EJW. 2018. A disturbed auxin signaling affects adventitious root outgrowth in *Solanum dulcamara* under complete submergence. Journal of Plant Physiology 224, 11–18.

Yang ZB, Liu G, Liu J, Zhang B, Meng W, Müller B, Hayashi Ki, Zhang X, Zhao Z, De Smet I. 2017. Synergistic action of auxin and cytokinin mediates aluminum◻induced root growth inhibition in Arabidopsis. EMBO reports 18, 1213–1230.

Yu M, Liu K, Liu S, Chen H, Zhou L, Liu Y. 2017. Effect of exogenous IAA on tension wood formation by facilitating polar auxin transport and cellulose biosynthesis in hybrid poplar (*Populus deltoids*× *Populus nigra*) wood. Holzforschung 71, 179–188.

Zažímalová E, Křeček P, Skůpa P, Hoyerova K, Petrášek J. 2007. Polar transport of the plant hormone auxin–the role of PIN-FORMED (PIN) proteins. Cellular and Molecular Life Sciences 64, 1621–1637.

